## Supplementary Information for "Subtype-specific single β1 integrin mechanics for activation, mechanotransduction and cytoskeleton remodeling"

Jo, Li et al.

### Contents

Supplementary Fig. 1-12

Supplementary Table 1

Legends for Movie 1-5

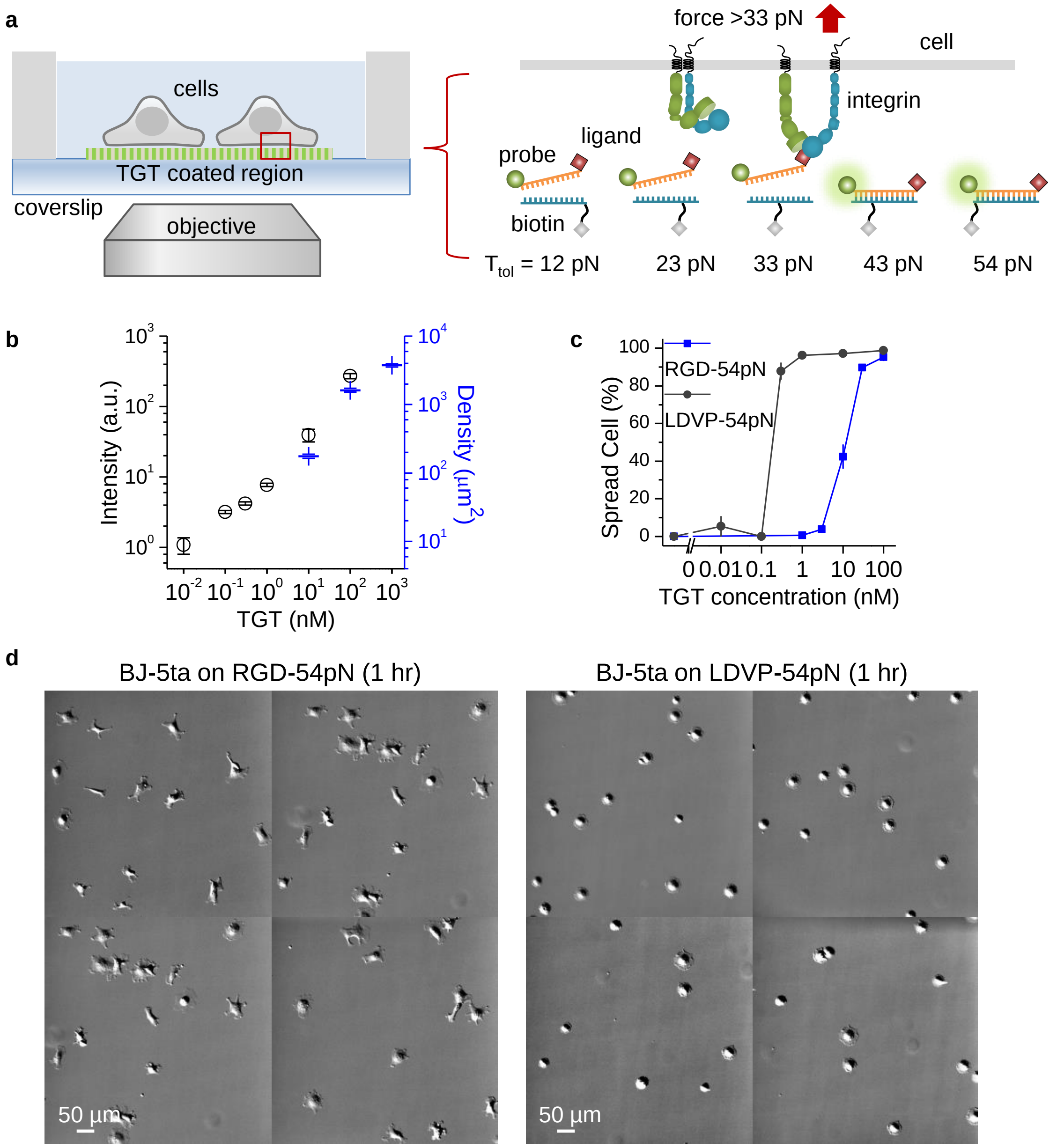


**Supplementary Fig. 1. Cells spreading on LDVP or RGD-TGT surface**

(a) Schematic of TGT with five different tension tolerances. (b) Fluorescent signal and immobilized TGT density according to TGT concentration on neutravidin coated PEG surfaces (10 min incubation). Fluorescence signals (Cy3) were measured using epifluorescence microscopy (black circles). The linearly increasing intensity shows that biotin-binding sites on the neutravidin coated surface are not saturated up to 100 nM. The data represent the mean ± SD from nine measurements (87 by 87 μm each). The molecular density was measured based on single-molecule detection (blue bars). See Supplementary Fig. 4a. TGTs with and without Cy3 were mixed for 100 nM (10% Cy3-TGT) and 1 μM (1% Cy3-TGT) conditions to use the identical imaging condition with 10 nM (100% Cy3-TGT) condition. The data represent the mean ± SD from five regions (87 by 87 μm each). (c) Percentage of spread cells on RGD-54pN or LDVP-54pN according to TGT concentration. TGT 100 nM was used across this study when the concentration is not specified. (d) Representative differential interference contrast (DIC) images (10X) of cells seeded on RGD-54pN (left) or LDVP-54pN (right) for 1 hr. Scale bar, 50 μm.

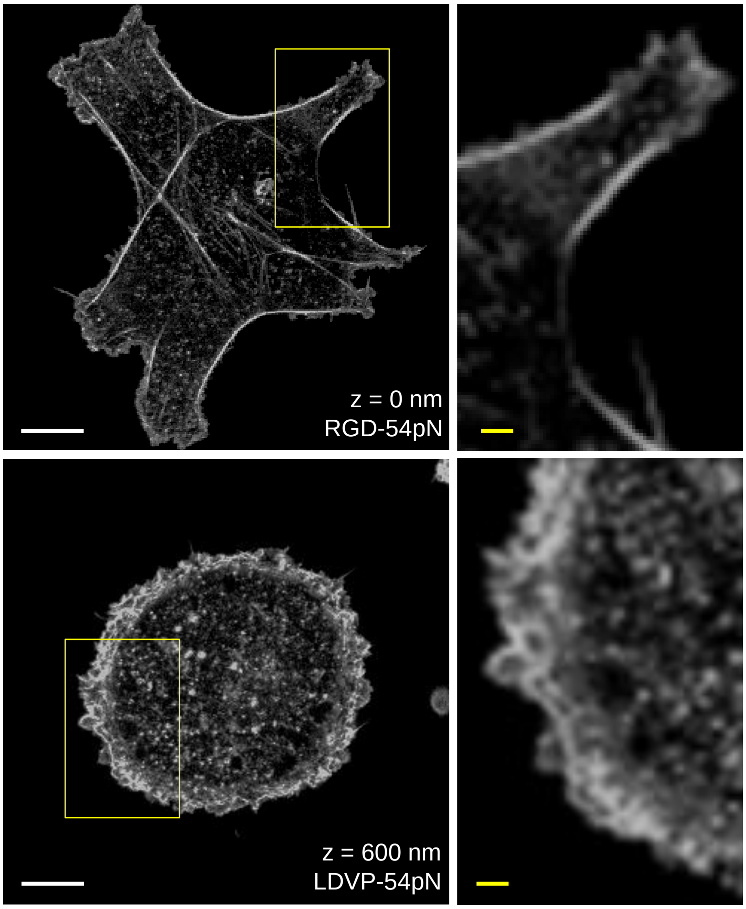


**Supplementary Fig. 2. Confocal images of cells spreading on LDVP or RGD-TGT**
Confocal images of phalloidin (AF555) show cytoskeletal structures of spreading cells. BJ-5ta fibroblasts were fixed after 1 hr spreading on RGD-54pN (top) and LDVP-54pN (bottom). Scale bars: 10 µm (white), 2 µm (yellow).

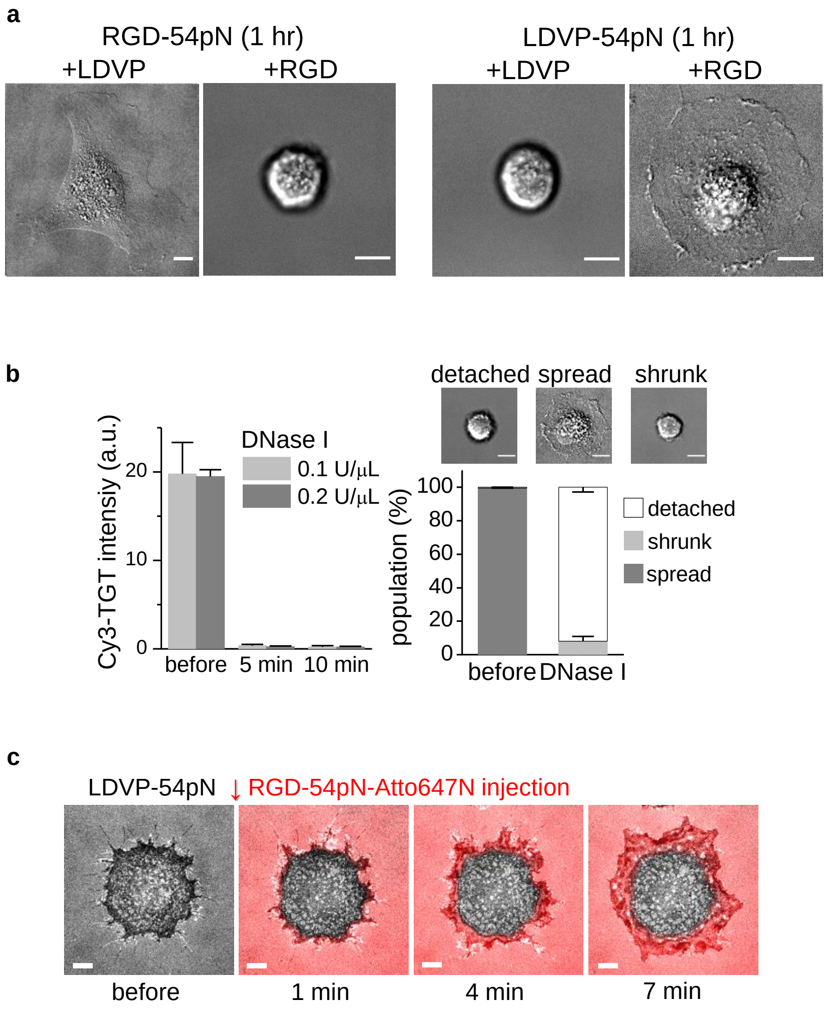


**Supplementary Fig. 3. RGD or LDVP-TGT mediated cell spreading**

(a) DIC images (60X) of cells seeded on TGT surfaces in presence of soluble LDVP (100 µM) or RGD (100 µM) for 1 hr. Scale bars, 10 µm. (b) To show that cell spreading is supported by TGT, DNase I was injected to the chamber with cells spreading on LDVP-54pN. Cy3 intensity of TGT surface shows the TGT density before and after DNase I treatment (left panel). The data represent the mean ± SD from three independent experiments. The percentage of cell shape before and 5 min after DNase I treatment (right panel). Cells were efficiently detached from the surface or shrunk with small contact. The data represent the mean ± SE from three independent experiments. (c) Additional immobilization of RGD-54pN. Cells were seeded on LDVP-54pN-Cy3 for 1 hr and RGD-54pN-Atto647N was injected (100 nM). The merged image is the overlay of RICM and Atto647N signal (red, TIRFM) on the surface. Scale bar, 10 µm.

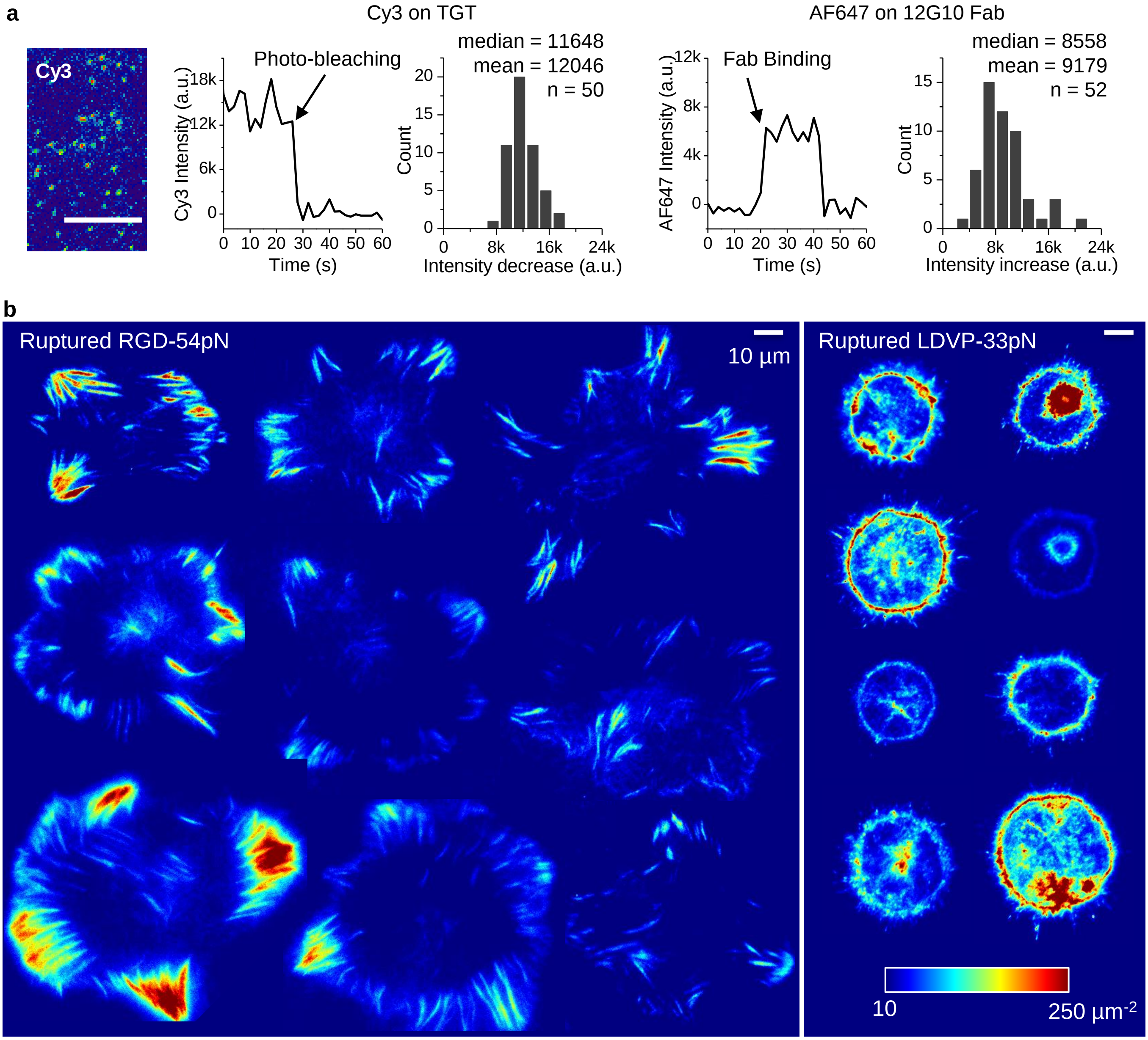


**Supplementary Fig. 4. High force transmission events recorded by qTGT.**

(a) Calibration of fluorescence signal for TGT, ruptured qTGT and bound Fab density. Fluorescent intensity was converted to the probe density by analyzing stepwise change of single-molecular signals. TIRF imaging was done for the region whose TGT or qTGT density is low enough to distinguish single-molecules (left image) on the same coverslip. The identical imaging condition was used except the exposure time (2 s for single-molecule imaging, 0.1-0.5 s for data acquisition). Scale bar, 10 µm. A time trace of Cy3 signal intensity with a photobleaching event is shown. a.u., arbitrary unit. The distribution of Cy3 signal from a calibration is shown. The median was used for TGT density calibration. For fluorescently labeled Fab (12G10 Fab, AF647), non-specific transient Fab binding events to a surface were monitored and analyzed in the similar way. The mean of the stepwise signal increase was used for the calibration considering multiple dye conjugated molecules (mostly 1-2 dyes). 40 or more events were collected for each calibration. This calibration was done for each experiment.. (b) Calibrated ruptured qTGT density maps ($\mu m^{-2}$) for a gallery of representative cells seeded on RGD-54pN (left) or LDVP-54pN (right). Scale bar, 10 µm.

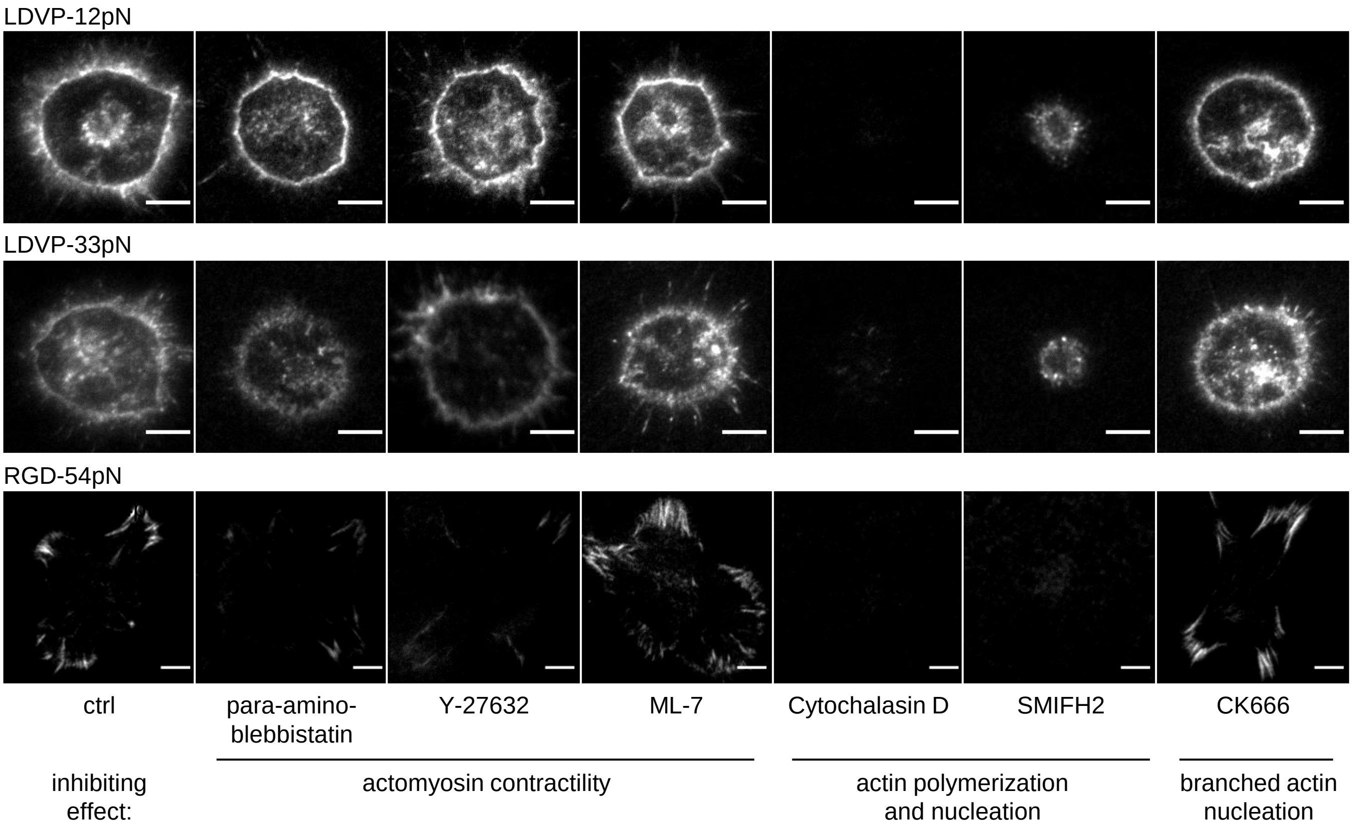


**Supplementary Fig. 5. Cytoskeletal inhibitor effects on TGT rupture**

Representative images of LDVP-12pN, LDVP-33pN and RGD-54pN (BHQ2-Cy3) ruptured by cells seeded for 1 hr in presence of cytoskeletal inhibitor (5 min preincubation): para-amino-blebbistatin (50 uM), Y-27632 (10 μM), ML-7 (10 μM), Cytochalasin D (10 μM), SMIFH2 (50 μM), and CK666 (50 μM). The outermost rupture signals reflect the spread cell area. Cells did spread when Cytochalasin D or SMIFH2 was added. Scale bar, 10 μm

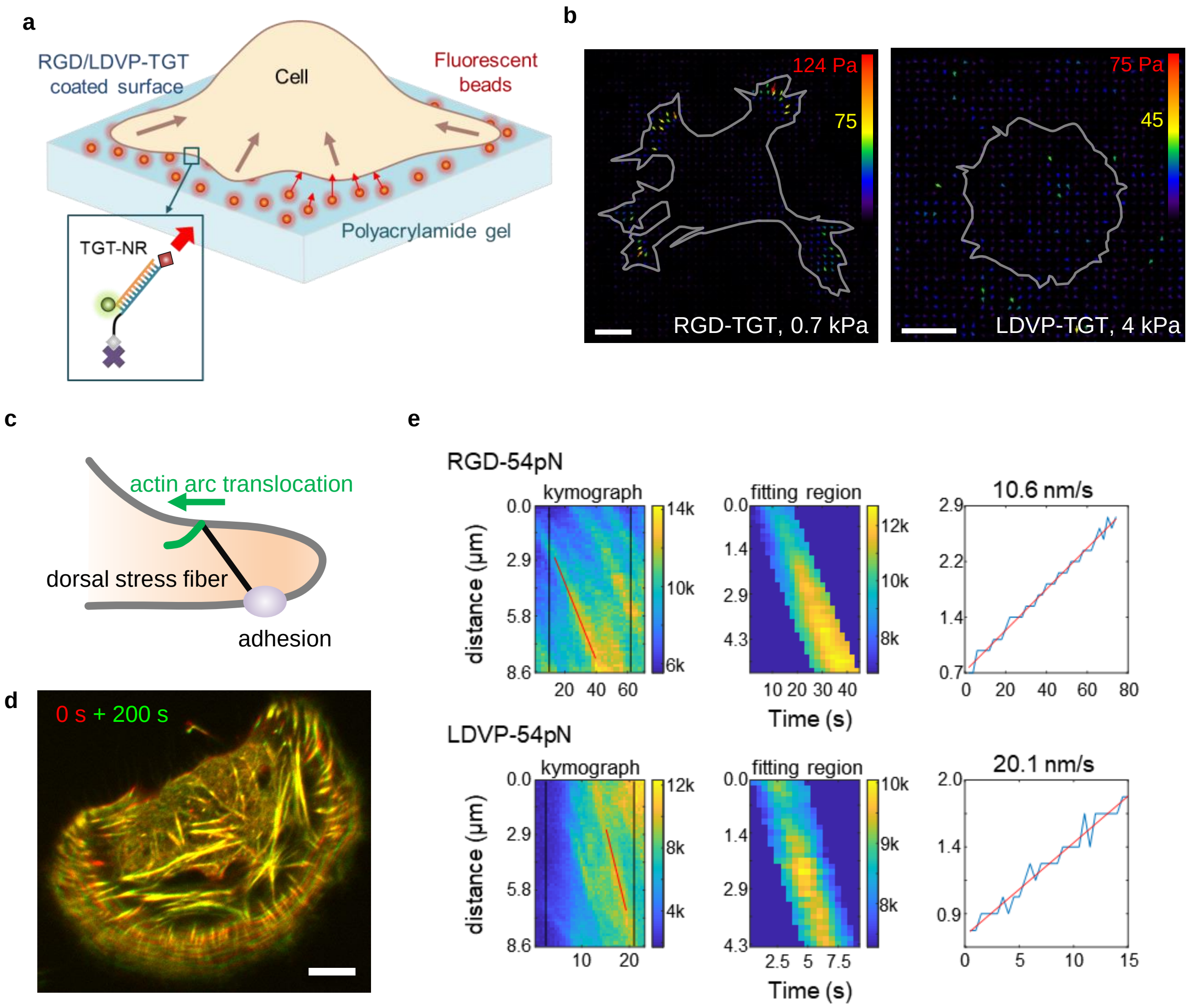


**Supplementary Fig. 6. Traction force and retrograde flow rate measurement.**
(a-b) Traction force of BJ-5ta fibroblasts on RGD or LDVP. (a) Schematic of traction microscopy using non-rupturable, TGT-like DNA linkers (Supplementary table 1). Ligands were conjugated on the biotinylated strands to prevent detachment of ligands from the surface. (b) Representative mechanical stress maps (force/area). Gray lines indicate the cell outlines. Scale bar, 10 μm. (c-e) Flow speed of actin arc was measured analyzing time-lapse TIRFM images of Sir-actin (10-20 nM). (c) Schematic of actin arc translocation connected to the adhesion. (d) Two time points were overlaid for a cell on RGD-54pN with pseudo-color to show the actin movement over time. Scale bar, 10 μm. (e) Representative kymographs for actin arc translocation analysis. The red lines indicate the spatiotemporal regions for linear fitting.

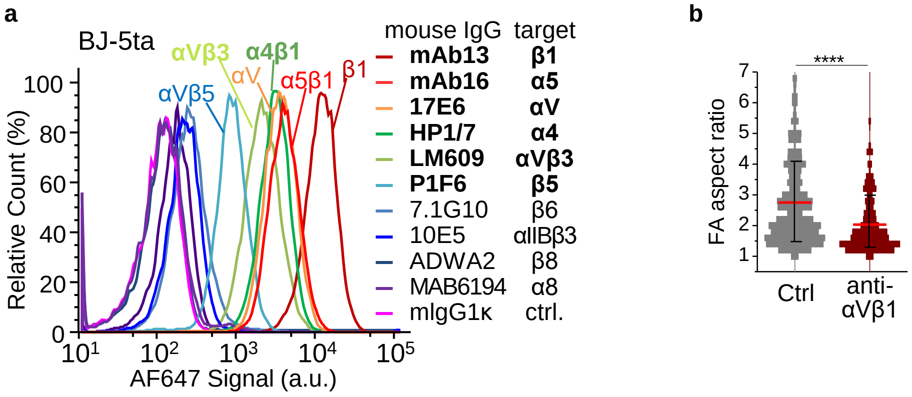


**Supplementary Fig. 7. Expression of integrins on BJ-5ta fibroblasts**

(a) Immuno-fluorescent flow cytometry of RGD-binding integrins and integrin α4β1 on BJ-ta cells using indicated mouse IgGs and AF647-labeled goat anti-mouse IgG. Antibodies with positive labeling are named in bold on right. (b) Aspect ratio of focal adhesions of cells seeded on RGD-54pN for 1 hr in absence or presence of inhibitory Biogen-αVβ1.5 Fab to αVβ1. Paxillin was immunostained (AF488) as the focal adhesion marker (clusters >1.2 $\mu m^2$; n = 2116 and 2251 from 13 and 17 cells). Two-sample t-test for p-values. ****$p<0.0001$.

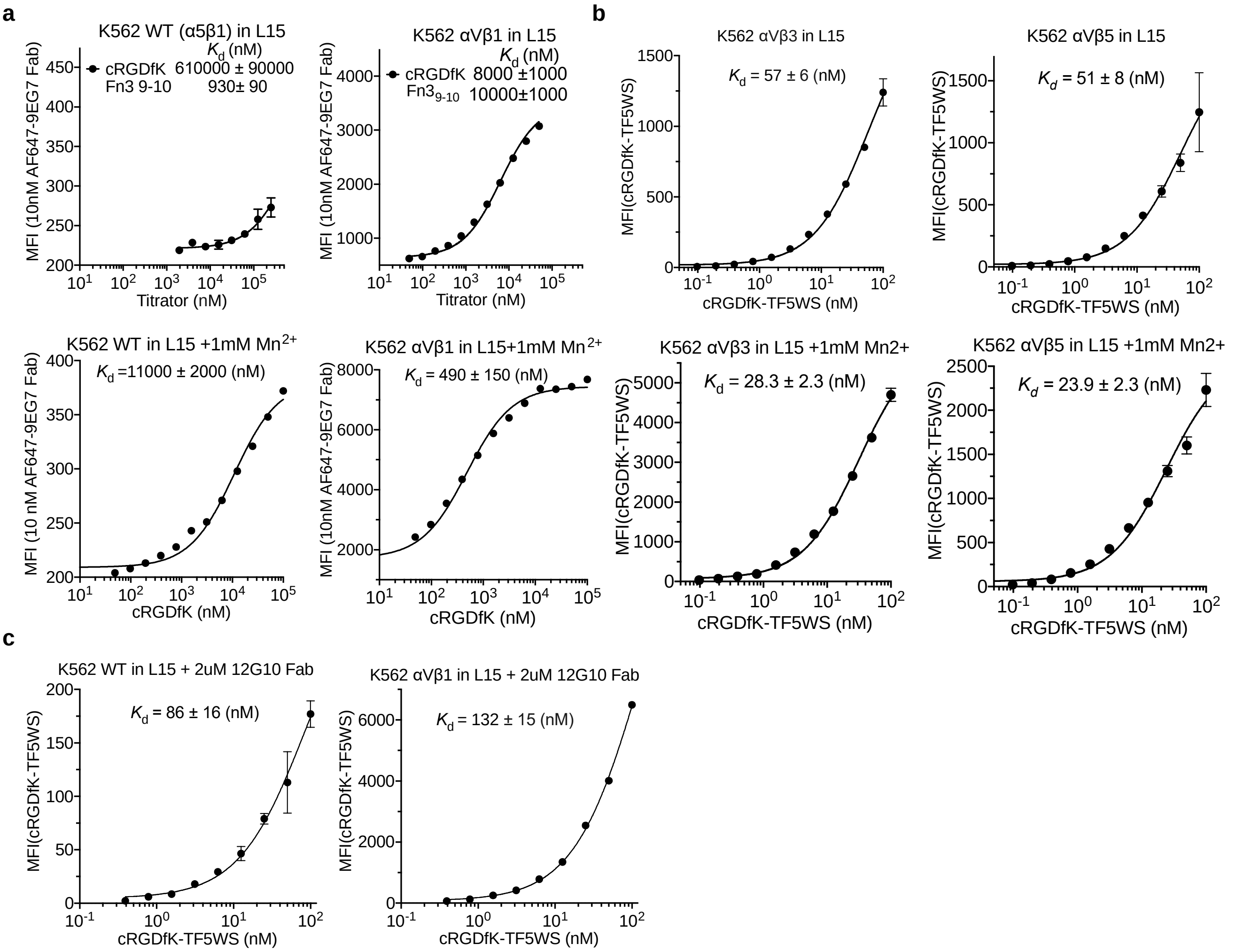


**Supplementary Fig. 8. Binding affinity of cRGDfK for RGD-binding integrins on cell surfaces.**
Affinities of cRGDfK peptide to intact integrins on K562 cells in L15 medium containing 1% BSA were quantified by flow cytometry without washing. (a) Affinity of cRGDfK to α5β1 on native K562 cells and αVβ1 on K562 stable transfectants was measured by enhancement of binding of AF647-9EG7 Fab (10 nM), which is specific for the β1 extended conformation. Mean fluorescence intensity (MFI) at each cRGDfK peptide concentration was fitted to three parameter dose-response curve for background MFI, maximum MFI, and the EC50 value which represents the $K_d$ value of cRGDfK peptide to the respective integrin. In L15 without $Mn^{2+}$, macromolecule fragment Fn3 9-10 was also included as a titrator to better determine the maximum MFI. Fits for cRGDfK and Fn3 9-10 shared the same background MFI and maximum MFI as fitting parameters. Binding to the natively expressed α5β1 in αVβ1 transfectants was negligible compared to αVβ1, owing to the much higher expression of αVβ1, as shown by the much lower affinity for fibronectin and much higher affinity for cRGDfK in the αVβ1 transfectants compared to native K562 cells in absence of $Mn^{2+}$ (upper panels), as well as the much higher affinity for cRGDfK in the αVβ1 transfectants compared to native K562 cells in presence of $Mn^{2+}$ (lower panels). (b) Affinity of cRGDfK to intact αVβ3 and αVβ5 was measured on K562 αVβ3 and K562 αVβ5 stable transfectants, respectively, by saturation binding of cRGDfK peptide with lysine side chain conjugated with TideFluor5WS (TF5WS) (custom synthesized by BACHEM). Background was measured with 10 mM EDTA present in the binding buffers. Background-subtracted MFI at each cRGDfK-TF5WS concentration was fitted to three parameter dose-response curve for background MFI, maximum MFI at saturation binding level, and $K_d$ value of cRGDfK-TF5WS to integrin αVβ3 or αVβ5. Binding to the natively expressed α5β1 in αVβ3 and αVβ5 transfectants was negligible, owing to the much higher expression of αVβ3 and αVβ5, respectively on the transfectants, and non-detectable specific MFI of cRGDfK-TF5WS (100nM ) on native K562 cells. (c) Intrinsic affinity of cRGDfK-TF5WS for EO state of integrin α5β1 or αVβ1 stabilized by 2 μM 12G10 Fab was measured using same methods as in (b).

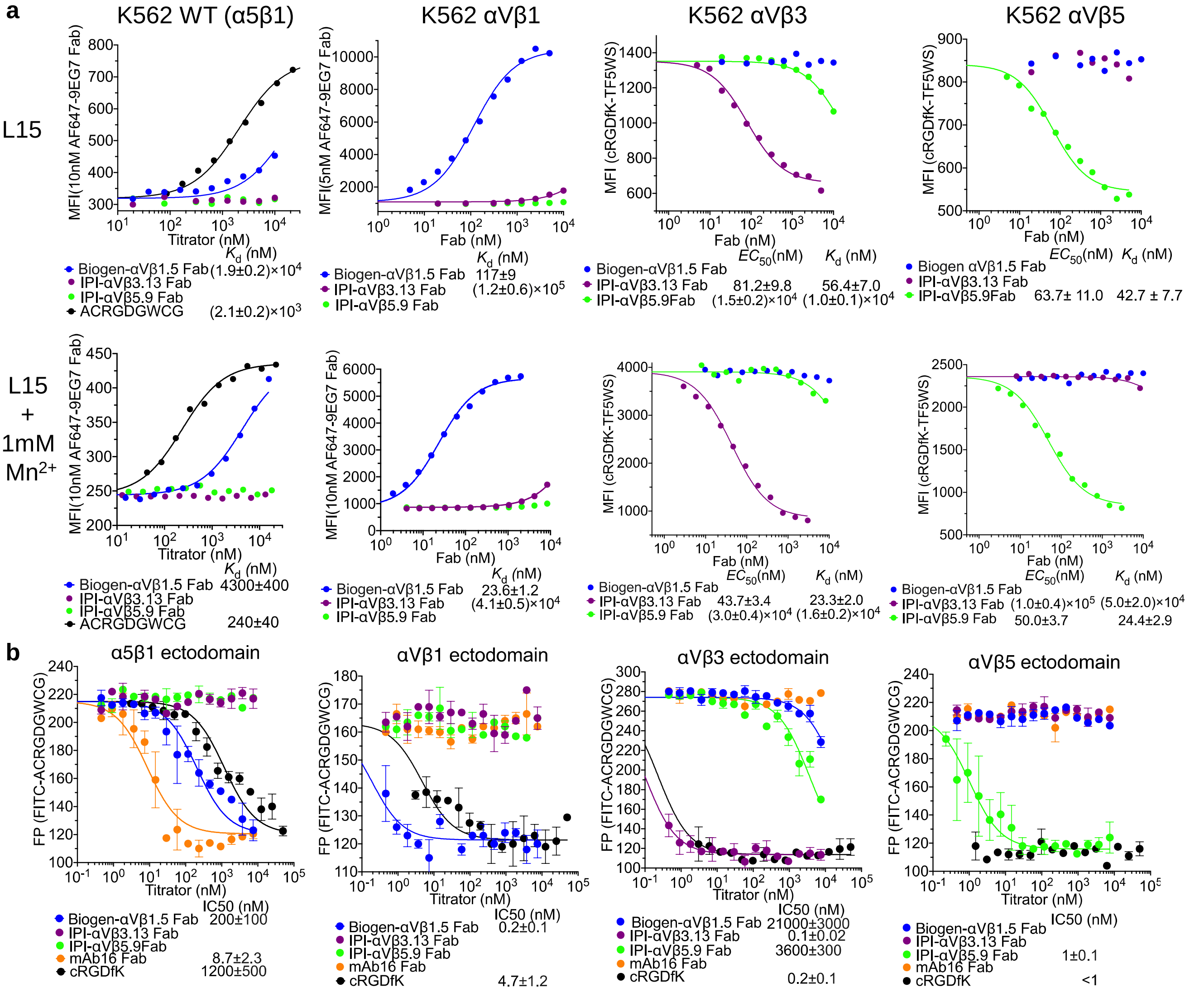


**Supplementary Fig. 9. Affinity and specificity of integrin inhibiting Fabs.**

(a) Affinities of RGD-mimetic Fabs to intact integrins on K562 cells in L15 medium containing 1% BSA with or without 1mM $Mn^{2+}$, quantified by flow cytometry without washing. Fab affinities for α5β1 expressed on K562 wild type cells and for αVβ1 expressed on K562 αVβ1 stable transfectants were quantified by enhancement of 10 nM AF647-9EG7 Fab binding. For measurement on α5β1, its high affinity binding peptide, cyclic-ACRGDGWCG, was also included as a titrator to better determine maximum MFI when all α5β1 is in the ligand bound EO state. MFI at each concentration of different titrators was globally fitted to three parameter dose- response curve, with background MFI and maximum MFI as shared fitting parameters, and $K_d$ value for each titrator as individual fitting parameter. Affinity of Fabs to integrin αVβ3 and αVβ5 were determined by competing with 25nM cRGDfK-TF5WS binding on K562 αVβ3 and K562 αVβ5 stable transfectants, respectively. MFI of cRGDfK-TF5WS at each concentration of different competitors were globally fitted to three parameter dose-response curve, with maximum MFI in absence of competitor and minimum background MFI as shared fitting parameters, and EC50 value for each competitor as individual fitting parameter. With the fitted EC50 value, $K_d$ of each competitor was calculated as $K_d = EC50 / (1+ C_L/K_{d,L})$, where $C_L$ is the concentration of cRGDfK-TF5WS used (25 nM), and $K_{d,L}$ is the binding affinity of cRGDfK-TF5WS to the respective integrin determined in Supplementary Fig. 8 under corresponding condition. (b) Specificity of inhibiting Fabs used in this study checked on soluble integrin ectodomains. IC50s of Fabs were quantified by competing 10 nM FITC-cyclic-ACRGDGWCG peptide binding to 50 nM α5β1, 200 nM αVβ1, 20 nM αVβ3 and 20 nM αVβ5 ectodomains, respectively, by fluorescence polarization (FP) assays. FITC-cyclic-ACRGDGWCG was labeled with FITC at the amino group of the same 6-aminohexanoic acid spacer used for TGT DNA strand attachment. Curves in each panel were globally fitted with three parameter dose response curve, with maximum FP value in absence of titrator and minimum FP value when all the integrin in solution is bound by titrator as shared fitting parameters, and IC50 as individual fitting parameter for each titrator.

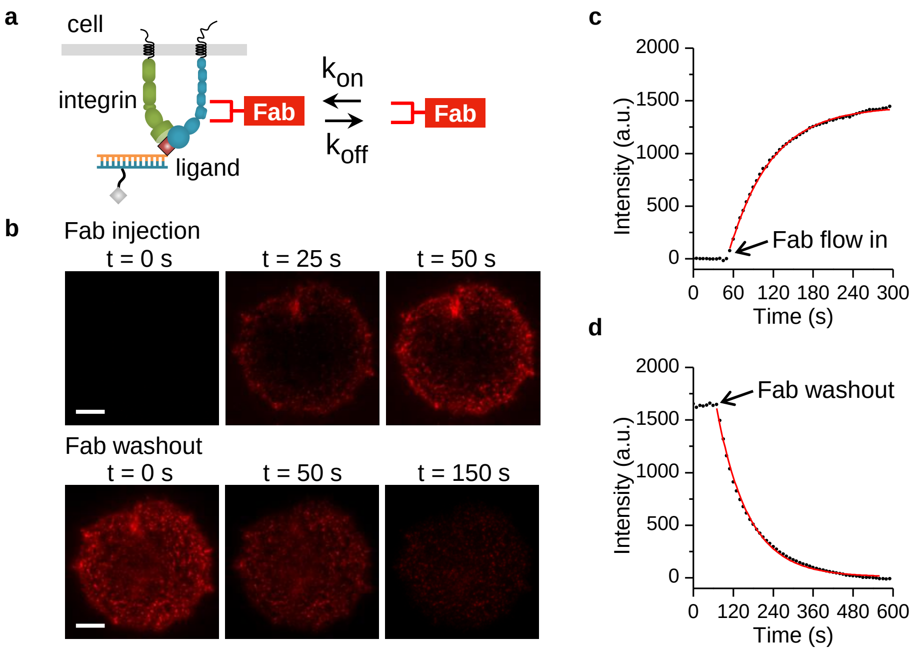


**e**

| Target | Fab | [Fab] | Cell | TGT | $k_{obs}$ ($s^{-1}$) | $k_{off}$ ($s^{-1}$) | $k_{on}$ ($\mu M^{-1}$ $s^{-1}$) | $K_D$ (nM) |
|---|---|---|---|---|---|---|---|---|
| β1E | 9EG7-AF647 | 20 nM | BJ-5ta | LDVP-54pN | 0.0146 ± 0.0006 | 0.0094 ± 0.0003 | 0.257 ± 0.019 | 37.3 ± 2.0 |
| β1EO | 12G10-AF647 | 20 nM | BJ-5ta | RGD-54pN | 0.0043 ± 0.0004 | 0.0030 ± 0.0003 | 0.066 ± 0.010 | 50.6 ± 8.3 |
| α4 | HP1/7-AF488 | 100 nM | BJ-5ta | LDVP-54pN | 0.0130 ± 0.0007 | 0.0050 ± 0.0002 | 0.080 ± 0.007 | 64.8 ± 7.3 |
| αv | 13C2-AF488 | 20 nM | BJ-5ta | RGD-54pN | 0.0099 ± 0.0005 | 0.0049 ± 0.0005 | 0.250 ± 0.015 | 20.3 ± 3.1 |

**Supplementary Fig. 10. Fab binding kinetics.**
(a) Kinetics were measured with Fabs specific to integrin α-subunits or conformational states of the β1 subunit. To check whether Fab binding kinetics were fast enough for live cell imaging, binding and dissociation rates of Fab were measured on cells spreading on RGD-54pN or LDVP-54pN. Cells were seeded in a flow chamber connected to a syringe pump. Fluorescent Fab solution was injected to measure binding kinetics and subsequently washed out to measure dissociation using TIRFM. (b) Representative images show the binding and dissociation of AF647-labeled 9EG7 Fab. Scale bar, 10 µm. (c-d) Representative association (c) and dissociation (d) curves of 9EG7 Fab on a single cell. The single-exponential fitting values were used for $k_{obs}$ and $k_{off}$. (e) Measured and calculated binding kinetic parameters. $k_{obs}$ = (binding time)$^{-1}$. $k_{off}$ = (dissociation time)$^{-1}$. $k_{on}$ = ($k_{obs}$ - $k_{off}$)/[Fab]. [Fab] is the concentration of Fab injected. $K_D$ = $k_{on}$/$k_{off}$. Values are mean ± SE of six measurements (two measurements from three cells).

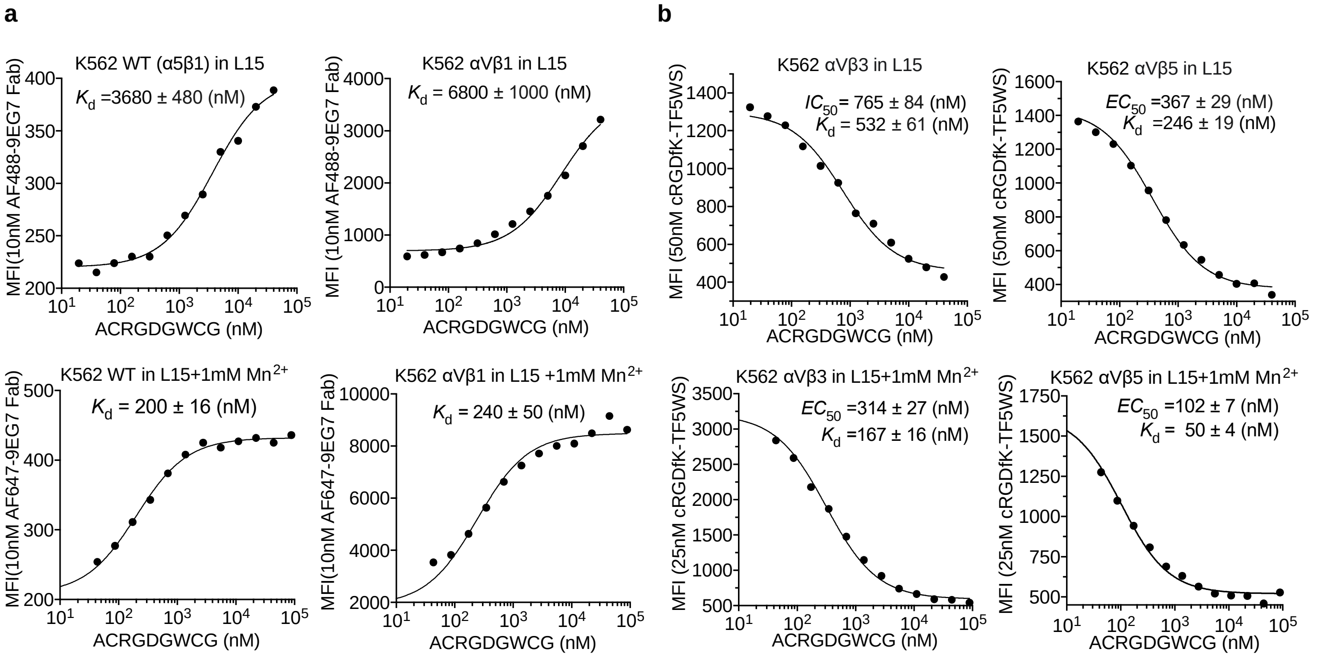


**Supplementary Fig. 11. Binding of cyclic-ACRGDGWCGK for RGD-binding integrins on cell surfaces.** Affinities of cyclic-ACRGDGWCGK peptide (ACRGD) to intact integrins on K562 cells in L15 medium containing 1% BSA were quantified by flow cytometry without washing. (a) Affinity to α5β1 on native K562 cells and αVβ1 on K562 stable transfectants was measured by enhancement of binding of AF647-9EG7 Fab (10 nM), which is specific for the β1 extended conformation. Mean fluorescence intensity (MFI) at each ACRGD concentration was fitted to three parameter dose-response curve for background MFI, maximum MFI, and the EC50 value which represents the $K_d$ value of ACRGD to the respective integrin. Binding to the natively expressed α5β1 in αVβ1 transfectants was negligible compared to αVβ1, owing to the much higher expression of αVβ1 on the stable transfectants, as also evidenced by the ~10-fold lower binding affinity of Fn3 9-10 to αVβ1 transfectants than native K562 cells (Supplementary Fig. 8a). (b) Affinity of ACRGD to intact αVβ3 and αVβ5, were measured on K562 αVβ3 and K562 αVβ5 stable transfectants, respectively, by competing with cRGDfK-TF5WS peptide binding . MFI of cRGDfK-TF5WS at each concentration of competitor, ACRGD, was fitted to three parameter dose-response curve, with maximum MFI in absence of competitor, minimum background MFI, and EC50 value for competitor as fitting parameters. With the fitted EC50 value, $K_d$ of ACRGD was calculated as $K_d = EC50 / (1+ C_L/K_{d,\,L})$, where $C_L$ is the concentration of cRGDfK-TF5WS used, and $K_{d,\,L}$ is the binding affinity of cRGDfK-TF5WS to the respective integrin determined in Supplementary Fig. 8 under corresponding condition. Contribution of natively expressed α5β1 to the MFI of cRGDfK-TF5WS on αVβ3 or αVβ5 transfectants were negligible, owing to the much higher expression level of αVβ3 or αVβ5 in the respective stable transfectants, and the much lower binding affinity of cRGDfK peptide for α5β1 (Supplementary Fig. 8). Thus, competitive binding monitored by the MFI of cRGDfK-TF5WS on αVβ3 and αVβ5 transfectants measures the binding of competitor ACRGD to αVβ3 and αVβ5, respectively.

### cRGDfK

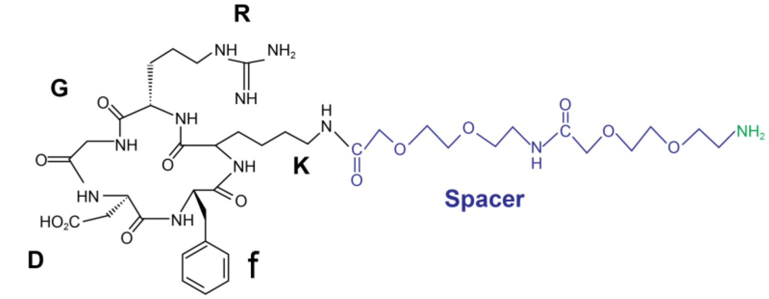


### MUPA-LDVPAAK

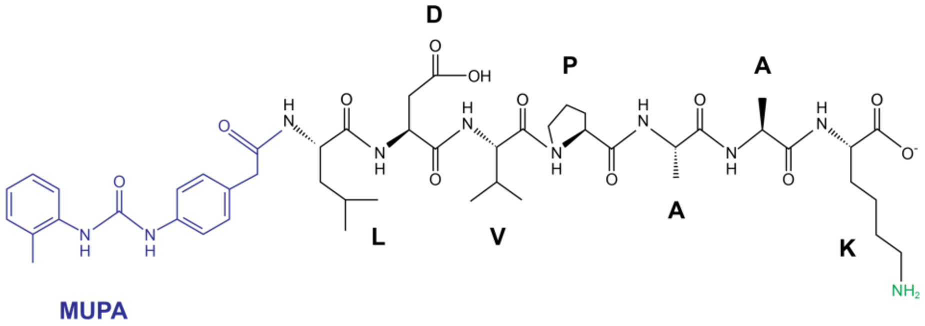


### Cyclic-ACRGDGWCGK

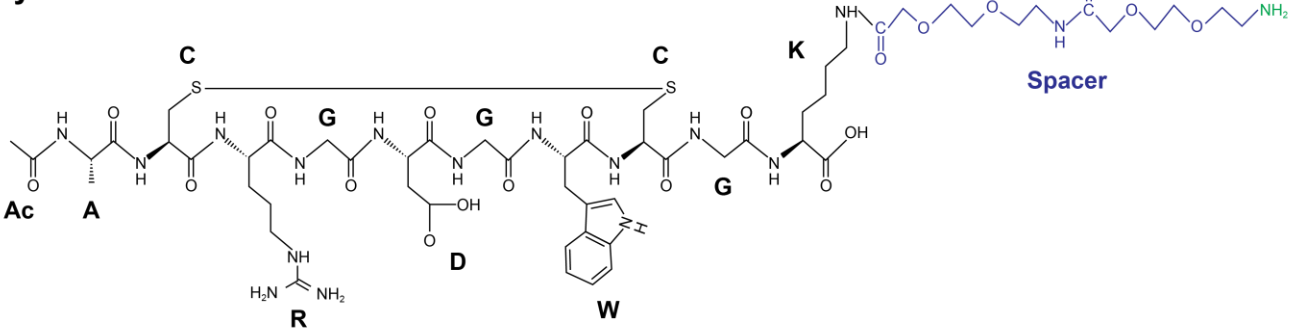


**Supplementary Fig. 12. Structural Formulas of peptidomimetic ligands used for TGTs.**
Chemical modifications of peptides and DNA attachment sites are colored in blue and green, respectively.

| Description | DNA sequence (5’ to 3’) |
|---|---|
| • Ligand strands for TGT (dye labeled) | |
| Cy3-labeled | /5Cy3/GGC CCG CAG CGA CCA CCC/3ThioMC3-D/ |
| for Atto647N-labeling | /5AmMC6/GGC CCG CAG CGA CCA CCC/3ThioMC3-D/ |
| • Ligand strand for quenched TGT | |
| RGD and LDVP-TGT | /BHQ-2/GGC CCG CAG CGA CCA CCC/Thiol C6 SS/ |
| ACRGD-TGT | /BHQ-2/GGC CCG CAG CGA CCA CCC/Amino C7/ |
| • Immobilization strands for TGT (not labeled) | |
| 12 pN | /5AmMC6/GGG TGG TCG CTG CGG GCC |
| 23 pN | GGG /iAmMC6T/GG TCG CTG CGG GCC |
| 33 pN | GGG TGG /iAmMC6T/CG CTG CGG GCC |
| 43 pN | GGG TGG TCG C/iAmMC6T/G CGG GCC |
| 54-pN | GGG TGG TCG CTG CGG GCC/3AmMO/ |
| • Immobilization strands for quenched TGT (dye labeled): | |
| 12 pN | /5Biosg/TTT TTT GGG TGG TCG CTG CGG GCC/3Cy3Sp/ |
| 23 pN | GGG /iAmMC6T/GG TCG CTG CGG GCC/3Cy3Sp/ |
| 33 pN | GGG TGG /iAmMC6T/CG CTG CGG GCC/3Cy3Sp/ |
| 43 pN | GGG TGG TCG C/iAmMC6T/G CGG GCC/3Cy3Sp/ |
| 54-pN | GGG TGG TCG CTG CGG GCC /iAmMC6T/TT TTT/3Bio/ |
| • Immobilization strands for non-rupturable TGT-like linker (used for traction force microscopy): | |
| Non-rupturable | /5AmMC6/GG GTG GTC GCT GCG GGC C/3ThioMC3-D/ |

/ThioMC3-D/ or /Thiol C6 SS/, disulfide modifier with C3 or C6 linker (for ligand conjugation);
/Amino C7/, amino modification with C7 linker (for ligand conjugation);
/5AmMC6/ or /3AmMO/ amino modification (for biotinylation);
/iAmMC6T/, internal amino modifier with C6 linker on thymine (for biotinylation);
/5Biosg/ or /3Bio/, biotin modification;
/BHQ-2/, Black Hole Quencher 2 modification;
/5Cy3/ or /3Cy3Sp/, Cy3 dye modification.

**Supplementary Table 1. DNA oligos for TGT synthesis**

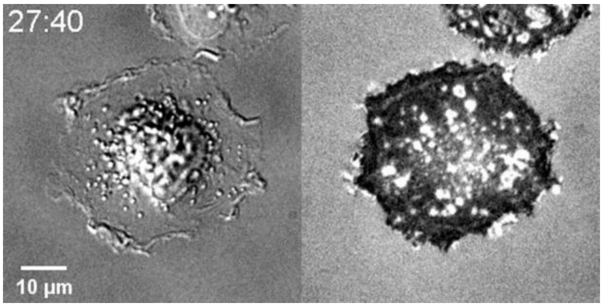


**Movie 1. Cell adhesion and spreading on LDVP-54pN TGT surface.**
BJ-5ta fibroblasts were seeded on LDVP-54pN coated surface. Cells promptly adhere and spread on the surface. DIC (left) and RICM (right) time-lapse images were taken at 20 s intervals. Scale bar, 10 µm.

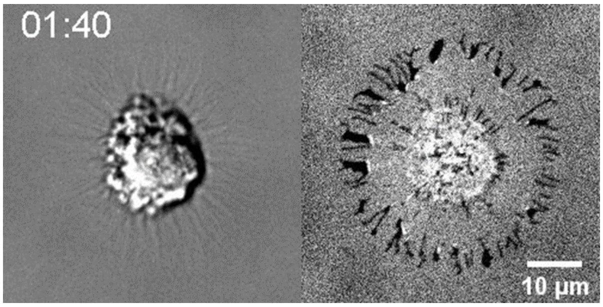


**Movie 2. DNase I-induced cell detachment from the TGT surface.**
To show that cell spreading is mediated by TGT, DNase I was added (time = 0 s) to the cells spreading on the LDVP-54pN surface. As TGTs were digested, cells lost their mechanical support and started to detach from the surface. DIC (left) and RICM (right) time-lapse images were taken at 20 s intervals. Scale, 10 µm.

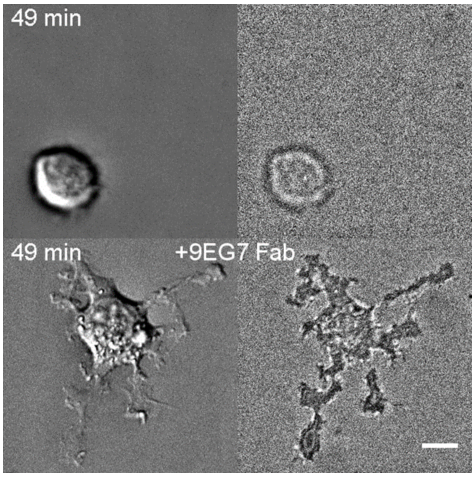


**Movie 3. Cells require >33pN tensile force on RGD-bound integrin to fully spread.**
BJ-5ta fibroblasts were seeded with and without 9EG7 Fab on RGD-33pN coated surface. Cells transiently adhered but failed to spread and detach from the RGD-33pN surface (top). On the same substrate with 9EG7 Fab (bottom), some cells adhered to the surface and started to spread, but spreading was limited compared with spreading on RGD-43pN and RGD-54pN surfaces. DIC (left) and RICM (right) time-lapse images were taken at 1 min intervals. Scale bar, 10 µm.

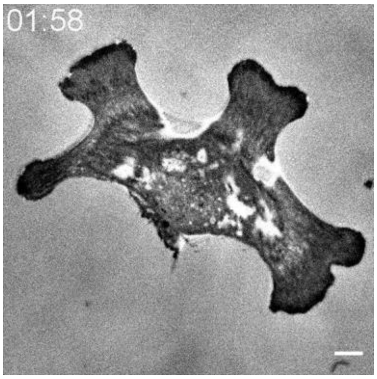


**Movie 4. Spreading though RGD-bound integrins activates elongation of cells initially spread through integrin α4β1 on LDVP.**
BJ-5ta fibroblasts were seeded on LDVP-54pN surfaces for 1 hr and RGD-54pN was injected (time = 0 min) into the cell chamber. Cells promptly start to spread further and get elongated. RICM time-lapse images were taken at 2 min intervals. Scale bar, 10 µm.

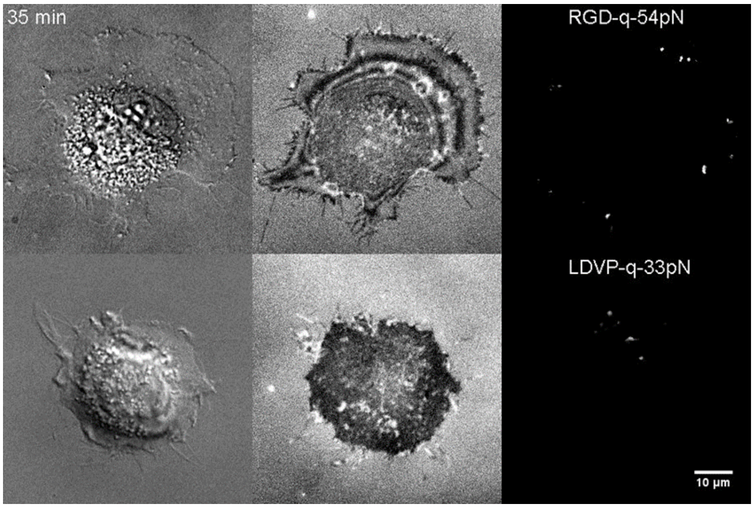


**Movie 5. Real-time molecular tension during cell spreading.**

BJ-5ta fibroblasts were seeded on RGD-54pN (top) and LDVP-33pN (bottom) BHQ2-Cy3 TGT. Real-time molecular force maps were obtained using qTGT rupture differential analysis (right). DIC (left) and RICM (middle) time-lapse images are also shown. Time-lapse images were taken at 1 min intervals for 40 min. Scale bar, 10 µm.
